# Proteomic analysis of liver tissue reveals *Aeromonas hydrophila* infection mediated modulation of host metabolic pathways in *Labeo rohita*

**DOI:** 10.1101/2021.11.16.468918

**Authors:** Mehar Un Nissa, Nevil Pinto, Biplab Ghosh, Urvi Singh, Mukunda Goswami, Sanjeeva Srivastava

**Affiliations:** Department of Biosciences and Bioengineering, Indian Institute of Technology Bombay, Powai, Mumbai 400076, India; Central Institute of Fisheries Education, Indian Council of Agricultural Research, Versova, Mumbai, Maharashtra 400061; Regional Centre for Biotechnology, Faridabad, 121001, India; Department of Biochemistry, Sri Venkateswara College, University of Delhi, India

**Keywords:** *Aeromonas hydrophila*, Liver proteomics, reprogramming, mass spectrometry, rohu

## Abstract

*Aeromonas hydrophila (Ah)* is an opportunistic Gram-negative bacterium and a serious global pathogen causing Motile Aeromonas Septicaemia (MAS) in fish and many other vertebrates. The pathogenesis of aeromonas septicaemia is complex and involves multiple perturbed pathways. Molecular analysis of host tissues could be a powerful approach to identify mechanistic and diagnostic immune signatures of disease. We performed a deep proteomic analysis of *Labeo rohita* liver tissue to examine changes in the host proteome during *Ah* infection. A total of 2525 proteins were identified of which 158 were found differentially expressed during *Ah* infection. Functional analysis of significant proteins identified the dysregulation of several metabolic enzymes, antioxidative proteins, cytoskeletal proteins and immune related proteins. Proteomic analysis revealed the alterations in the cellular defence mechanisms including phagolysosomal killing and apoptosis during *Ah* infection. Our systemic approach revealed the protein dynamics in the host cells to explore the putative biological processes underlying the metabolic reprogramming of the host cells during *Ah* infection. Our findings paved the way for future research into the role of Toll-like receptors (Tlr3), C-type lectins (Clec4e) and metabolic enzymes in *Ah* pathogenesis leading towards host directed immunotherapies to tackle the *Ah* infection in fish.

**IMPORTANCE:** Bacterial disease is one of the most serious problems in aquaculture industry. *Aeromonas hydrophila* (*Ah*), a Gram-negative bacterium causes motile aeromonas septicaemia (MAS) in fish. Small molecules that target the metabolism of the host have recently emerged as potential treatment possibilities in infectious diseases. However, the ability to develop new therapies is hampered due to lack of knowledge about pathogenesis mechanisms and host-pathogen interactions. Molecular level analysis of host tissues could be helpful in finding mechanistic immunological markers of diseases. We examined alterations in the host proteome during *Ah* infection in *Labeo rohita* liver tissue to find cellular proteins and processes affected by *Ah* infection. Our systemic approach revealed protein dynamics underlying the host cells’ metabolic reprogramming during *Ah* infection. Our work is an important step towards leveraging host metabolism in targeting the disease by providing a bigger picture on proteome pathology correlation during *Ah* infection.

## INTRODUCTION

The intensification of aquaculture system leads to the occurrence of diseases that reduce the quality of fish and fishery products and in turn cause economic loss. Aquaculture is experiencing a massive loss of production due to a variety of reasons of which more than 50% are due to diseases especially in developing countries (1). Aeromonads have been recognised as the most common bacterial pathogens responsible for bacterial fish diseases. Aeromonads include *Aeromonas hydrophila (Ah)*, *A. veronii, A. sobria,* and *A. caviae* which harm a wide range of hosts (2). Among these, *Ah* is highly infective and results in economic loss at alarming levels (3). *Ah* is a gram-negative opportunistic pathogen. Infectious signs of dropsy, hemorrhages, ulcers and necrosis have been observed in all stages of carps and other freshwater fish species like *Arapaima gigas*, Nile tilapia (*Oreochromis niloticus*) and catfish (4–7).

Proteomic analysis has become crucial for understanding the fundamental mechanisms of bacterial resistance and virulence. This has resulted in a greater understanding of pathogen biology and interaction with host that can be well addressed through holistic approaches like proteomics (8). Proteomics approaches have been utilised to explore the antibiotic resistance mechanism in *Ah.* With the help of proteomic approaches, it has been reported that the quinolones resistance in *Ah* might involve increased expression of SOS response-related proteins while decreasing those of chemotaxis (9). Further, the proteomic analysis of carp intestinal mucosa revealed that the differentially expressed proteins belong to MHC II protein complex and immune response, throwing light on the metabolic processes that are important during *Ah* infection (10). In *Ah* infected Wuchang bream (*Megalobrama amblycephala*), proteomic analysis of hepatopancreas revealed that *Ah* infection affected antioxidative proteins through complex regulatory mechanisms and decreased immunological ability (11). In the gills of Zebrafish, mucosal immune response was found enhanced during *Ah* infection (12).

Fish majorly depend on their innate immune system as their first line of defense which is an important factor in disease resistance (13). *Ah* infections cause changes in the host metabolism, but the mechanisms that determine the nature and severity of these changes are still not known completely. In order to explore the infection mediated metabolic changes in the host, we performed proteomic analysis of liver tissue of *Ah* infected *Labeo rohita. L. rohita* is the most important aquaculture species among the three Indian major carp species in polyculture system. Liver is a metabolically active tissue and has been recognized as a central immunological organ with high exposure to circulating antigens and endotoxins from the gut microbiota, particularly enriched for innate immune cells (14). This study focussed on proteomic characterisation of infected liver to understand *Ah* pathogenesis and to look into the physiological changes and host response to *Ah* infection.

## RESULTS

### Aeromonas hydrophila infection alters proteomic profile in host liver tissue

Discovery based proteomics data was acquired for eight samples (Details in Material and Methods) through high resolution mass spectrometry and a label-free quantitation (LFQ) approach was utilised for detection and quantification of the host proteins. The significantly altered proteins (p-value 0.05) were considered for the identification of the best panel of proteins to differentiate the control and *Ah* infected group. Further using a targeted mass spectrometry-based approach of selected/multiple reaction monitoring (SRM/MRM), selected protein targets were validated where data was acquired for eleven samples (five AH and six Control). Finally, we looked into how *Ah* (Ah) infection impacted the host’s physiological processes (Fig. 1).

**Figure 1.**
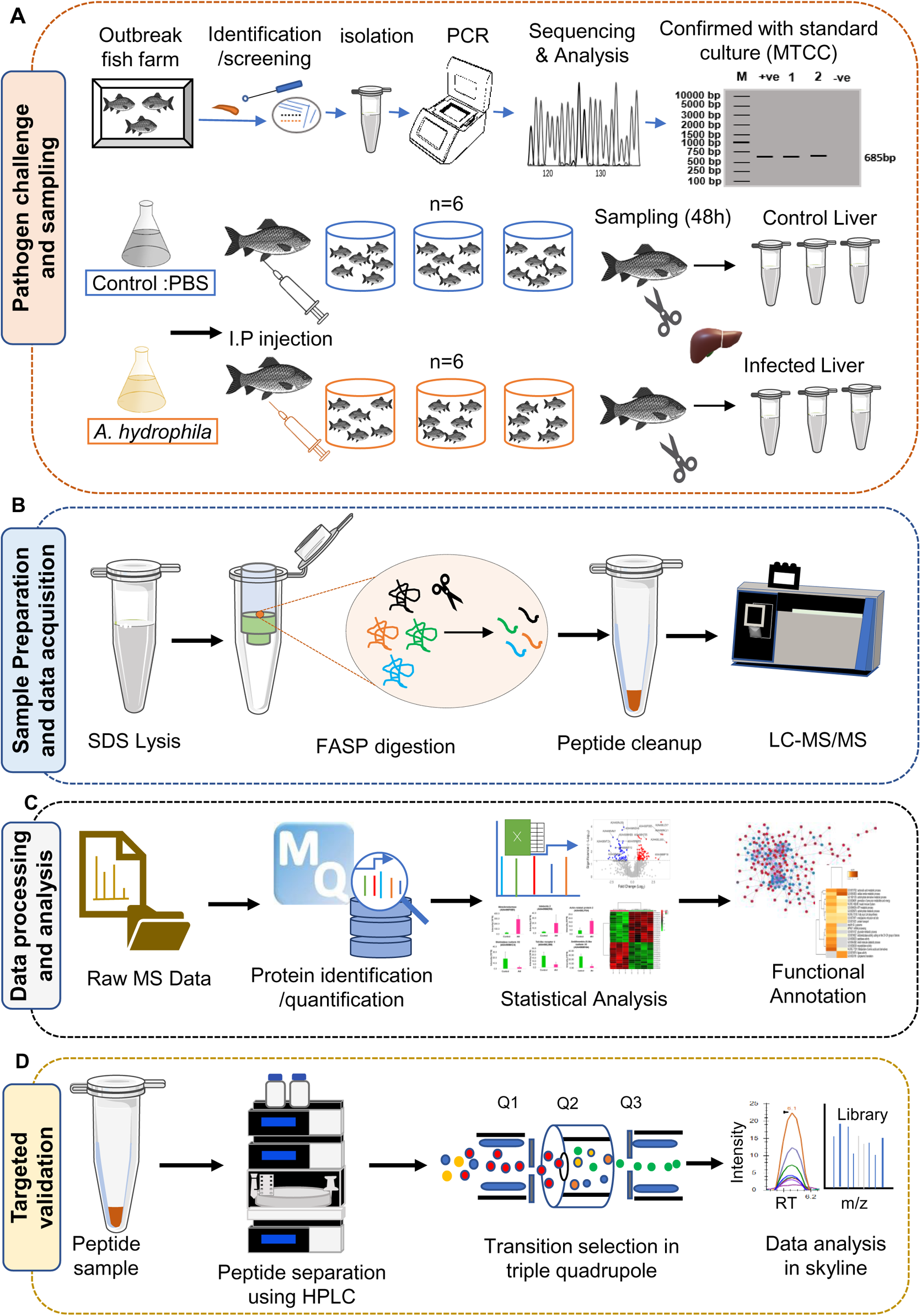
Overview of sample collection and experimental workflow: **(A)** Rohu collected from the outbreak farm was screened, kidney sample was streaked onto TSA plate; identical colonies were selected from TSA plate followed by DNA isolation, targeted Polymerase Chain Reaction (PCR) for 16sRNA gene and sequencing for identification of bacteria. To re-confirm MT374248 isolated strain, *A. hydrophila* from MTCC culture (1739T) was used. For challenge study, Phosphate Buffer saline (PBS) and *A. hydrophila* was injected intraperitoneally to Control and AH group (*Ah* challenged) respectively. After 48h of infection, fish were euthanized and liver samples were collected (Details in text). (**B)** For proteomics analysis, tissues were lysed in Sodium dodecyl sulphate (SDS) containing buffer followed by protein digestion using filter assisted sample preparation (FASP) method. Peptide samples were cleaned before subjecting to mass spectrometry for Data dependent acquisition (DDA). (**C)** Acquired raw mass spectrometry data (.raw) was analysed using MaxQuant software for identification and quantification of proteins. Statistical analysis was performed to identify the differentially expressed proteins followed by functional analysis. **(D)** Targeted proteomic validation of selected proteins using selected reaction monitoring was done where peptide sample was subjected to High performance liquid chromatography (HPLC) followed by target precursor and transition selection in Triple quadrupole mass spectrometer for acquisition of spectral data. SRM data was analysed using Skyline software where targeted data was compared with Spectral library prepared from DDA data.

Comparison of proteomic profiles from uninfected and infected liver tissue was performed to define changes to the host proteome during *Ah* infection. After analysing through label-free quantification approach, on an average ~1650 proteins were quantified in each replicate (Table S1) except one of the samples from infected group (AH1) which was removed from downstream analysis because of poor mass spectrometric results compared to other samples (Fig. S1A-B). Further statistical analysis was performed for 7 out of 8 samples; four of the control samples (Liv-C1 to Liv-C4) and three infected samples (Liv-AH2 to Liv-AH4). We observed 390 proteins to be enriched in control group (viz. quantified in at least 75% of control samples and not detected or detected in less than 33% samples of AH condition) and concurrently 113 proteins were enriched in AH group (*Ah* infected) which might be a reflection of different proteome expression under healthy and diseased condition (Fig. S1C, Table S1). Differential expression analysis was performed for 1157 overlapped proteins between the two groups. In comparison to the control group, we observed that in the *Ah* infected group, 59 proteins exhibited a decreased abundance while 98 proteins showed an increased abundance representing a total of 157 dysregulated proteins during *Ah* infection (Fig. 2A, Table S1) (see Materials and Methods for significance threshold values). We employed Partial Least Squares Discriminant Analysis (PLS-DA) to identify the proteins that can distinguish the AH group (*Ah* infected) from the Control group on the basis of variable importance in projection (VIP) Score. Top 30 features (proteins) that can clearly classify between two groups are represented (Fig. 2B). Unsupervised hierarchical clustering analysis clearly clustered the samples with similar abundance profiles and stratified the bacterial infected and control group as represented for top 30 features (Fig. 2C). Proteins that could differentiate the infected group form control group mapped to diverse molecular functions as described below. Comparative protein abundances for few of the differentially expressed proteins is represented in Fig. 2D including Metalloreductase STEAP4 (Steap4), Glutamine gamma-glutamyltransferase (Tgm1) and Actin-related protein 2 (Actr2) showing increased abundance and Biotinidase (Btd), Toll-like receptor 3 (Tlr3), Antithrombin-III-like isoform X1 (Serpinc1) proteins showing a decreased abundance as a result of *Ah* infection.

**Figure 2.**
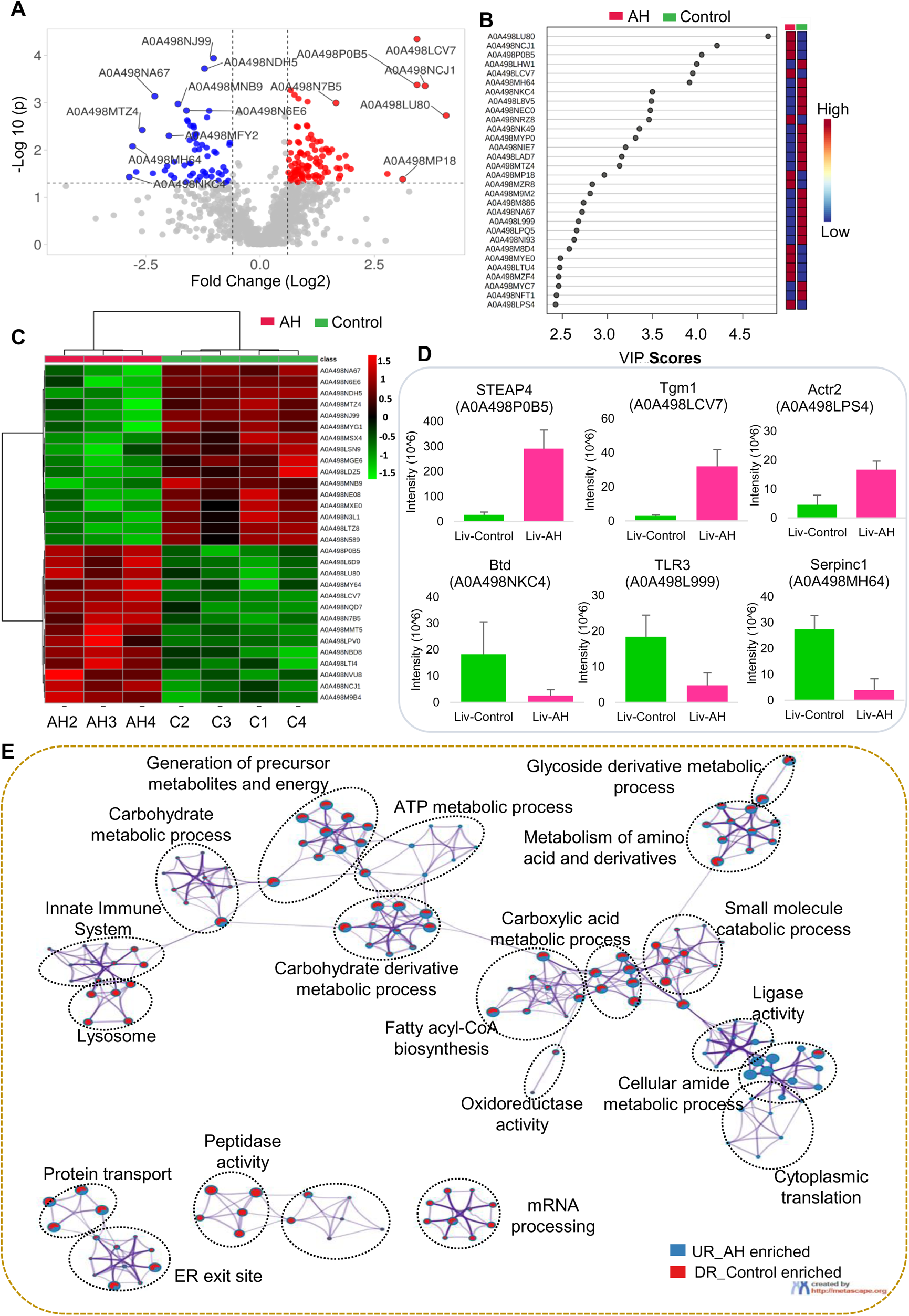
Shotgun proteomic analysis reveals altered host proteomic signatures in liver during *Ah* infection: **(A)** Volcano plot depicting significantly altered protein candidates (Fold Change 1.5, p ≤ 0.05), where red and blue colours represent upregulated and downregulated proteins respectively. (**B)** Top 30 altered key proteins found on the basis of VIP Score. **(C)** Heatmap showing differential expression (abundances) of top 30 significant proteins (Student t-test, p ≤ 0.05) across 4 replicates of control tissues (C-1, 2, 3, 4) and 3 replicates of *Ah* infected tissues (AH-2, 3, 4). **(D)** Bar graphs representing altered abundances of six functionally important proteins (three up and three downregulated) across Control and AH group. **(E)** Pathway enrichment network (using Metascape) showing functional annotation of protein coding genes. Each gene ontology (GO) and/ or pathway term is represented by a circle node, where its size is proportional to the number of input genes mapped to that GO term, and nodes present within specific circle belong to the same cluster. Terms with a similarity score > 0.3 are linked by an edge (the thickness of the edge represents the similarity score). Color codes for pie sector represents attributes of genes from two input lists i.e., red and blue colour represent number of gene candidates mapped from list 1 (DR_Control: downregulated significant and Control enriched proteins) and list 2 (UR_AH: upregulated significant and disease enriched protein set), respectively.

Differentially expressed proteins (DEPs) along with the proteins quantified only in one group (i.e., Control enriched or AH enriched) were considered for functional analysis (gene ontology) to further look into their biological association. Proteins that showed decreased abundance in AH group (*Ah* infected) belong to the processes like metabolism of amino acids, endopeptidase and peptidase activity, lysosomal processes, oxidoreductase activity and innate immune system. However, proteins with increased abundance during the infection were mainly mapped to carboxylic acid metabolism, cellular amide metabolism, Cytoplasmic ribosomal components, cytoplasmic translation and ligase activity (Fig. 2E, Fig. S2, Table S2).

### Dynamics of host metabolic pathways and protein-protein interactions during Ah infection

We performed protein-protein interaction (PPI) and pathway analysis for the DEPs and enriched proteins to see how the *Ah* infection affected the overall composition of the liver proteome. For the downregulated proteins, the enriched pathways and processes include the processes like lysosome, apoptosis, metabolism of xenobiotics by cytochrome P450, retinol metabolism, pantothenate metabolism, beta alanine metabolism, drug metabolism, metabolism of RNA and endocytosis (Fig. 3A-B). Pathway enrichment analysis of the upregulated proteins revealed the involvement of these proteins in several biological processes and pathways including innate immune system, signaling of B cell receptor, proteosome pathway, ribosome, carbon metabolism, protein synthesis (translation) and protein processing in ER (Fig. 3C-D). Dysregulation of these processes may indicate the severity of infection and its adverse effects on overall physiological processes.

**Figure 3.**
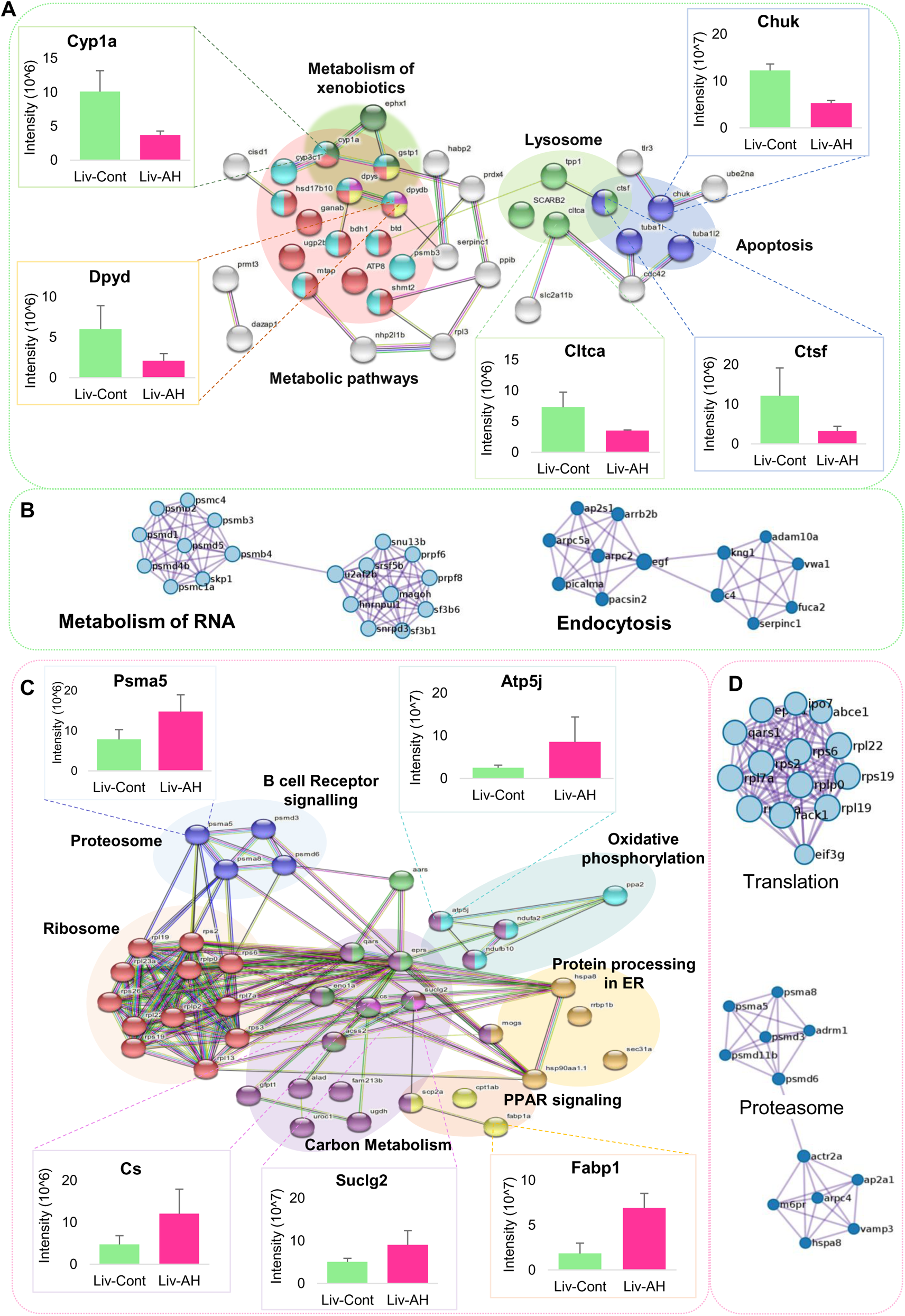
Landscape of altered pathways and interaction networks during *Ah* pathogenesis: **(A)** Interaction map depicting Protein-Protein interaction between significantly downregulated proteins and their involvement in altered pathways (based on STRING analysis). Bar plots showing differential protein abundance (intensity) for a few downregulated proteins (Cyp1a, Dpyd, Chuk, Cltc and Ctsf) from different pathways, based on label-free quantification (LFQ) **(B)** PPI enrichment analysis reveals top Molecular Complex Detection (MCODE) network components i.e., mRNA metabolism and Endocytosis among significant downregulated proteins and disease enriched proteins (based on Metascape analysis). **(C)** Protein-Protein interaction for significant upregulated proteins and their respective altered pathways during infections (STRING). Bar plots showing differential protein abundance (intensity) for a few upregulated proteins (Psma5, Atp5j, Cs, Suclg2 and Fabp (Fabp1) from different pathways, based on LFQ analysis. **(D)** PPI enrichment analysis reveals top MCODE network components i.e., Translation and Proteasome among significant upregulated proteins and disease enriched proteins (Metascape).

### *A. hydrophila* infection alters anti-oxidative defence system and hepatic xenobiotic metabolism

Infections are generally accompanied by a disturbance of the normal cellular homeostasis. Biotinidase protein (Btd) expression was found to be decreased by 7-folds during *Ah* infection (Fig. 2D, Table S1). Biotinidase recycles protein-bound biotin, resulting in the regeneration of free biotin which is essential for carbohydrate, protein and fat metabolism. Biotin deficiency has been reported to impair normal growth and immune functions (15). Another protein Antithrombin-III-like isoform X1 (Serpinc1) was also decreased by ~7 folds during *Ah* infection. Antithrombin is a natural coagulant synthesised in the liver and also involved in the anti-inflammatory signaling responses (16). Proteins like Glutathione transferase (Gst-A0A498NKS6), Glutathione peroxidase, Cytochrome P450 family (Cyp1a, Cyp2f2), Epoxide hydrolase (Ephx1), Dihydropyrimidinase (Dpys) and Dihydropyrimidine dehydrogenase [NADP (+)] (Dypd), involved in Xenobiotics and Drug metabolism (17) were downregulated during *Ah* infection. Proteins related to antioxidative system were also downregulated such as Peroxiredoxin-4 (Prdx4), Gst and Prenylcysteine oxidase 1 (Pcyox1). Downregulated proteins were also mapped to biological processes involved in general functions of liver such as Monooxygenase (Cypa1a) and Epidermal retinol dehydrogenase 2-like protein (Sdr16C5) from Retinol metabolism, Dihydropyrimidinase (dpys) and Dihydropyrimidine dehydrogenase (dpydb) from Pantothenate and CoA biosynthesis and beta-Alanine metabolism, Glycine hydroxymethyltransferase (Shmt2) which is involved in several biological processes like carboxylic acid metabolism, amino acid metabolim and energy metabolic processes (Fig. 3A, Table S2).

### *A. hydrophila* infection alters the cytoskeleton, lysosomal and apoptotic mechanism

Bacteria manipulate host cytoskeletal proteins for their mobility and distort the normal cytoskeleton. Few of the host cytoskeletal proteins including Actin-related protein 2 (Actr2), Alpha N-catenin (Ctnna2), Nucleolin (Ncl) were observed with an increased abundance change of 3.6-fold, 1.8-fold and 1.8-fold, respectively (Table S1). Under normal conditions, these proteins maintain the cytoskeletal homeostasis and the protein Actr2 is an ATP-binding component of the Arp2/3 complex that helps in cell motility by mediating the development of branching actin networks in the cytoplasm. The functioning of the Arp2/3 complex is required for vesicle trafficking, lamellipodium protrusion, and pathogen movement inside the host cell during infections. (18). Important proteins from lysosome pathway (KEGG ID: dre04142) showed a decreased trend in AH group (*Ah* infected). These include Cathepsin F-like protein (Ctsf, 3.7-fold decreased), Tripeptidyl-peptidase 1-like protein (Tpp1, 4.2-fold decreased), Lysosome membrane 2-like protein (Scarb2, 2.6-fold decreased) and Clathrin heavy chain (Cltca, 2.1-fold decreased). Different subunits of VATPase including Atp6voc, Atp6v1a, Atp8, Atp2b1 and palmitoyl-protein thioesterase 1 (Ppt1), Ras-related rab-15 (Rab15) and Ras-related Rab-1A (Rab1a) were also downregulated. Phospholipid scramblase 2-like (Plcsr3) which is a member of phospholipid scramblases, reported to mediate apoptosis by ATP independent bidirectional migration of phospholipids (19) has been observed with a 3.6-fold decreased abundance in *Ah* infected group (Table S1).

Another important protein, Glutathione-dependent dehydroascorbate reductase (Chuk) from apoptosis pathway was also found to be downregulated in AH group by 2.3-fold. Proteins from the immune system; Toll like receptor 3 (Tlr3) and C-type lectin domain family 4 member E-like protein (Clec4e/Cd207) and Chuk were found downregulated by 1.6, 1.9 and 2.3-fold, respectively, during the *Ah* infection. TLRs recognise pathogen-associated molecular patterns originating from microbes play a crucial function in macrophage maturation and activation and C-type lectins are also known to enhance adaptive immune responses (20).

### Proteomic alterations related to immune functions, protein synthesis and Carbon metabolism

As expected, we observed significant increase in the host proteins belonging to proteosome and ubiquitin dependent degradation pathways (KEGG Pathway ID: dre03050) and B cell receptor (BCR) signalling, (Reactome Pathway ID: DRE-1168372) (Fig 3B). Many biological activities rely on ubiquitination, including transcriptional regulation, cell cycle progression, signal transduction, protein transport, immunological responses and pathogenesis. Bacterial infections may take advantage of the host’s ubiquitin system to manipulate the immune response for their own benefits (21). Proteins like CCT-theta (Cct8), Intelectin 2 (Intl2), Proliferation-associated 2G4 (Pa2g4a), Galectin (Lgals3) and Unconventional myosin-Ic-like isoform X1 (Myo1c) which showed increased abundance in the *Ah* infected group, mapped to Innate immune system. Intelectin (Intl2) is a galectin binding lectin which has shown more than 6-fold increase during *Ah infection*. Intelectin has been shown to agglutinate bacteria, most likely due to its carbohydrate-binding ability, implying its role in innate immune system during the *Ah* infection (22). Also, the upregulated proteins such as subunits of proteasome; Psma8, Psma5, Psmd3, Psmd6, involved in downstream signaling events in BCR might be an indication of immune response during infection. Further, a panel of upregulated proteins mapped to ribosomal complex (viz. Rpl7a, Rps6, Rplp2, Rpl13), aminoacyl-tRNA biosynthesis (viz. Qars, Eprs, Aars), and protein processing in endoplasmic reticulum (viz. Hsp90aa1, Rrbp1, Hspa8, Sec31a, Mogs). The increase in protein synthesis machinery during infections could be a modulated host response to maintain protein homeostasis and to promote host cell immune response against pathogen (23).

Proteins of carbon metabolism like 6-Phosphofructo-2-kinase (Pfk2), 2-Phospho-D-glycerate hydro-lyase (Eno1a), Citrate synthase (Cs), Succinyl-ligase (Suclg2), Acetyl-coenzyme A synthetase (Acss2) and oxidative phosphorylation (OXPHOS) as ATP synthase-coupling factor 6 (Atp5j), Inorganic diphosphatase (Ppa2), Complex-I proteins (Ndufa2, Ndufb10) were found to be increased during *Ah* infection in this study. Few proteins from OXPHOS including ATP synthase protein 8 (Atp8) and NADH Dehydrogenase (Ndufa6, Ndufb6, Ndufa12) were found to be control enriched and were not quantified in infected group. Also, proteins from fatty acid metabolism and peroxisome proliferator-activated receptors (PPAR) signaling pathways (viz. Fabp, Scp2, Cpt1) were upregulated in the *Ah* infected group. Alterations in Scp2 expression have been linked to several other liver functions such as bile acid metabolism, biliary lipid secretion, hepatic cholesterol storage and synthesis (24). PPAR signaling plays active role in regulating inflammatory responses in innate and adaptive immunity. The anti-inflammatory and antibacterial characteristics of PPAR activation may benefit the host during bacterial infections (25). Mineral and oxidant homeostasis was also altered during the *Ah* infection as a few related proteins such as Metalloreductase Steap4 (Steap4) and Glutamine gamma-glutamyltransferase (gGT) were upregulated in the *Ah* infected group. Stress response proteins including Heat shock HSP 90-alpha (Hsp90aa1) and Heat shock cognate 71 kDa (Hspa8) were also upregulated during the infection.

### Targeted proteomic validation of dysregulated proteins using Selected Reaction Monitoring approach

The targeted proteomics data using SRM approach, was acquired for 27 differentially expressed proteins selected based on the discovery proteomics data (DDA followed by LFQ) (details in Methods section). The analysis ended up with a list of 328 transitions and 33 peptides belonging to sixteen proteins (plus 18 transitions for spiked in peptide) (Table S3). Nine proteins had at least two significant peptides. Group comparison analysis was performed in Skyline between 5 AH and 6 control samples which showed that the overall trend for these proteins is similar to that of DDA data in terms of increased or decreased abundance in *Ah* infected samples (AH condition) (AH condition) (Table S3). Among these, four proteins had three or more significant peptides passing the cut-off p value of 0.05 and fold change criterion of 1.5. These included three downregulated proteins viz. Bdh1 (D-beta-hydroxybutyrate mitochondrial), Chuk (D-Glutathione-dependent dehydroascorbate reductase) and Ugp2 (UTP-glucose-1-phosphate uridylyltransferase) (Fig. 4A-C) and one upregulated protein Atp5j (ATP synthase-coupling factor 6) (Fig. 4D). Individual sample wise peak area intensities for all the peptides of these proteins is represented in supplementary data (Fig. S3A-D, Table S3). Bdh1 is an important enzyme involved in lipid catabolism, Chuk protein has been reported for its role in apoptosis and Ugp2 is a carbon metabolic enzyme. Among the upregulated protein, the protein abundance changes were validated for Atp5j which is an important protein of oxidative phosphorylation. Additionally, five other proteins were found to be significant with 2 peptides only. These included three downregulated proteins namely, Dpys, Tpp1 and Sdr16c5, and two upregulated proteins ribosomal protein Rpl7a and Steap4 protein (Fig. S4).

**Figure 4.**
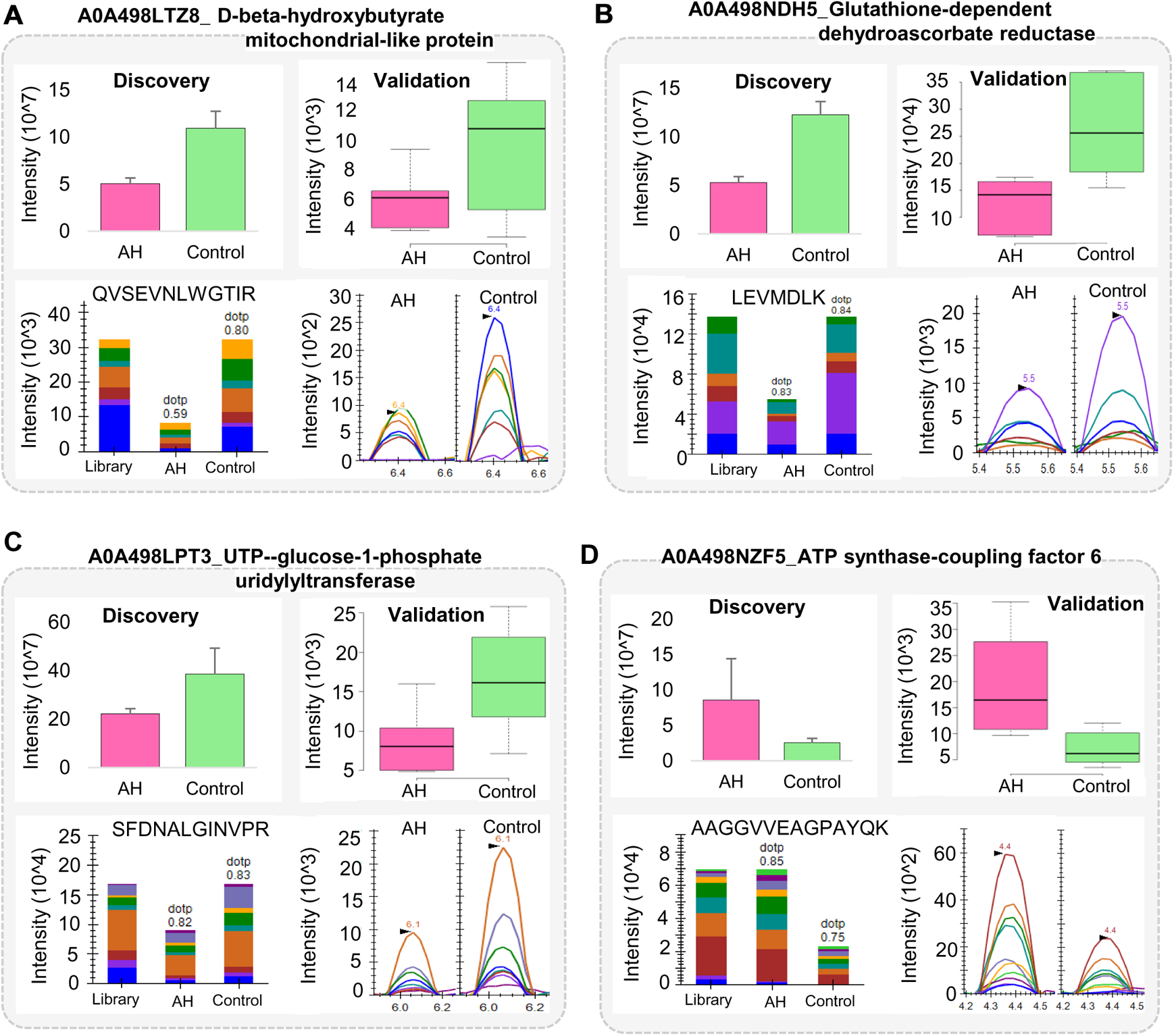
Experimental validation using targeted approach for changes in protein abundance during *Ah* infection: **(A-C)** Boxplots, peak area intensities and Peak shapes for downregulated proteins viz. Bdh1, Chuk and Ugp2, respectively and **(D)** Boxplot, peak area intensity and Peak shape for upregulated protein i.e., Atp5j in the AH condition (Ah infected group) compared to the control condition. In each figure (Left-Right), Upper panel represents the protein wise intensities based on the shotgun analysis (DDA abundance) and the targeted analysis (MRM intensity), respectively for each AH and Control condition showing similar trend in both the analyses. Lower panel shows the bar plots for peak area (with dot product (dotp) value based on match with spectral library) and spectral peak group comparison for a representative peptide of the same protein. Different colors in the bar plots and spectral peaks (in lower panel) represent different product ions of the same peptide.

## DISCUSSION

Bacterial diseases are the most prominent cause of mass mortality in freshwater aquaculture system. Among the bacterial diseases, the disease caused by Aeromonas group has great significance. Proteomic analysis of liver tissue of *Aeromonas hydrophila (Ah)* infected *L. rohita* was performed to explore the perturbed proteins and pathways in *Ah* pathogenesis. This is the first comprehensive proteomic analysis carried out for liver tissue of *Ah* infected *L. rohita*. Pathway enrichment analysis of the upregulated and downregulated proteins revealed the involvement of these proteins in several biological processes that could provide insights for the pathogenesis of *Ah* infection. Our results summarised the key proteins and pathways involved in metabolic reprogramming of host cells during *Ah* infection. Pathways like lysosome pathway, apoptosis, metabolism of xenobiotics by cytochrome P450, retinol metabolism, pantothenate metabolism, beta alanine metabolism and drug metabolism were found to be majorly mapped by downregulated proteins. However, innate immune system, signaling of B cell receptor, proteosome pathway, ribosome, carbon metabolism and protein processing in ER were mainly mapped to upregulated proteins.

Our results showed a decrease in Biotinidase enzyme in liver tissue during *Ah* infection. Biotinidase is an important enzyme responsible for regeneration of biotin. In the head kidney, spleen, and skin of grass carp (*Ctenopharyngodon idella*), biotin deficiency lowered the mRNA levels of anti-microbial compounds (hepcidin, mucin and defensin), increased the levels of pro-inflammatory cytokines as interferon γ2 (IFN-γ2), interleukin 6 (IL-6), IL-1, IL-8 and tumour necrosis factor α (TNF-α) (15). During *Ah* infection, Antithrombin showed a decrease in *Ah* infected group compared to the control group. Besides its function in coagulation, antithrombin has been reported to exhibit anti-inflammatory responses against many gram negative and positive bacteria as it showed affinity to bind with lipopolysaccharide (LPS) on bacterial surfaces (16).

We observed that *Ah* infection affects the antioxidative and xenobiotic potential of the host. Liver is recognised as the primary site of detoxification and xenobiotic metabolism. Decreased protein abundance of several proteins was observed including few CYP proteins (Cyp1a, Cyp2f2, Cyp2g1, Cyp3a) related to xenobiotic metabolism and antioxidation, during *Ah* infection. Dysregulation of CYP genes was observed in channel catfish liver, gills and intestine after infection with *Edwardsiella ictaluri* where transcripts of several members of Cyp2 were downregulated as in our case (26). Monooxygenase Cyp1a protein which is downregulated in our study, showed an increase at transcriptomic level in the kidney, gill, testis and liver of Nile Tilapia after *Ah* infection (27). Proteins related to oxidative mechanism were decreased in AH group including Peroxidoxin 4 (Prdx4), Glutathione transferase, Glutathione peroxidase and Prenylcysteine oxidase 1. Our results for Prdx4 match with the report for human cell line samples where a proteome level decrease was observed in Prdx4 along with Prdx2 and Prdx6 after Lymphocytic Choriomeningitis Virus infection (28). We found a significant increase in the abundance of Steap4, a metalloreductase enzyme involved in iron and copper homeostasis. It is reported to play important role in cell responses to inflammatory and oxidative stresses and its expression can be modulated by hypoxia and cytokines such as IL-1 beta and TNF-alpha (29). Our results for Steap4 are similar to those reported in spleen of rainbow trout in response to *Yersinia ruckeri* infection (30). Such observations indicated an overall effect of *Ah* infection on general body functions and growth of fish through oxidative stress or dysregulation of its antioxidant capacity.

Our results suggest that *Ah* infection tends to affect the process of phagolysosomal killing by downregulating several proteins involved in the process. Proteins like Cathepsins, Tpp1, Ppt1 are cysteine and serine proteases or hydrolases that may play role in pathogen killing (31, 32). Vatpase enzymes are important for maintaining the acidic pH inside the lysosomes whereas the other proteins (as Rab proteins) are important for cargo transportation inside the cell (33). Another protein, Phospholipid scramblase 2-like (Plcsr3) was observed decreased during *Ah* infection. Plcsr3 has been reported to assist in the recognition of apoptotic cells by macrophages (19). Alterations in these proteins could be the outcome of the pathogenic processes that favor intracellular survival of the pathogen in the host cells (Fig. 5A).

**Figure 5.**
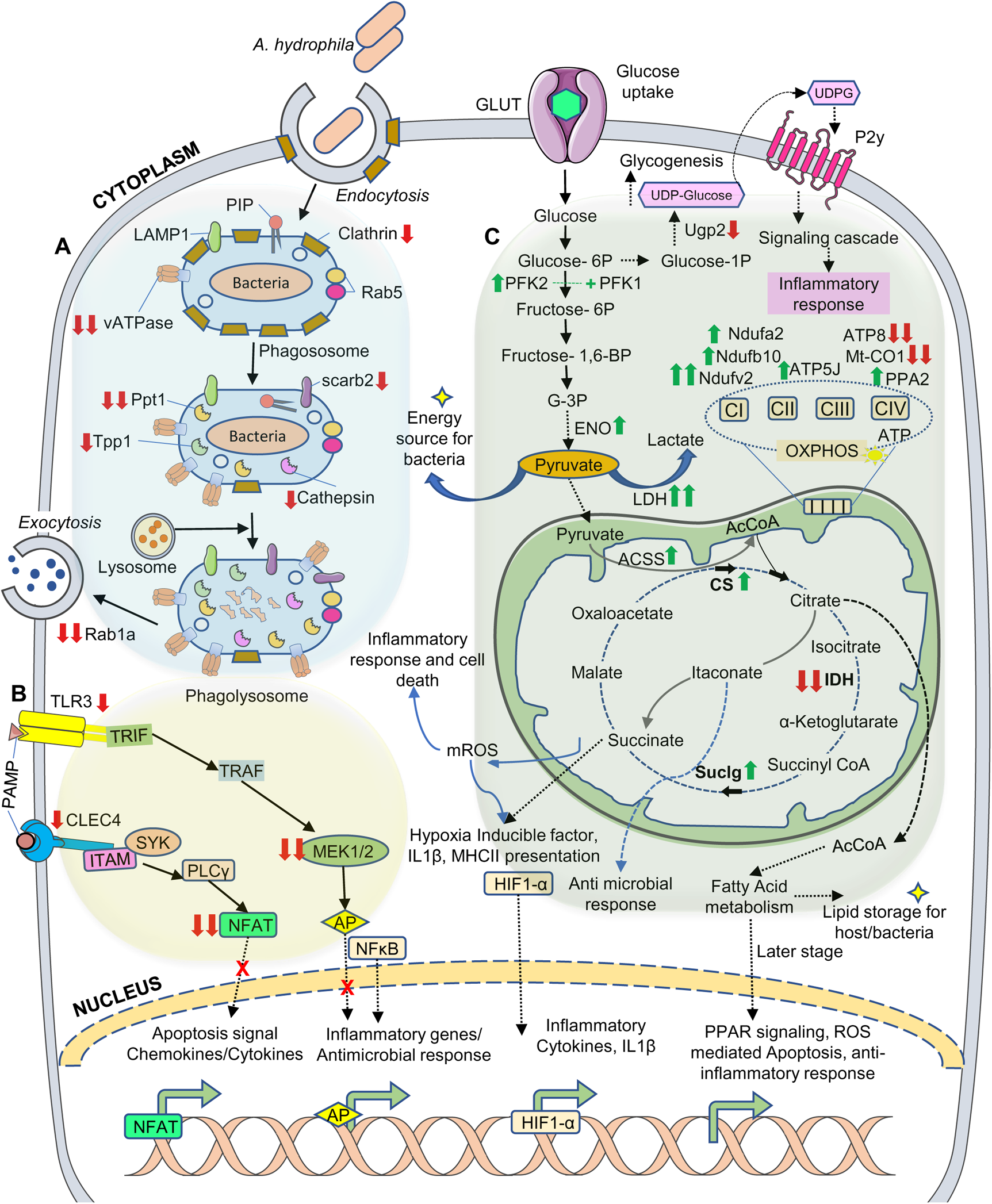
Schematic representation of interplay between host defence and pathogen survival during *Ah* infection in *L. rohita*: **(A)** Once the *Ah* bacterium (A. *hydrophila*) is engulfed by the host cell through phagosome, it undergoes a series of transformations by interacting with the subcompartments of the phagocytotic pathway and finally becomes phagolysosome after fusing with lysosome. During the process, the phagosome becomes highly acidic through proton pumping by VATPase, which is essential for the intracellular killing of microbe. Material that is not useful is exported through exocytosis. In the infected host cell, several proteins involved in phagocytosis-exocytosis pathway include Rab proteins, clathrin, lysosomal-associated membrane proteins (LAMP), lysosomal integral membrane protein (Scrab2), VATPase, hydrolases and proteases (cathepsin, Tpp1). During *Ah* infection, many of the host proteins are dysregulated, and this may help bacteria survive inside the host cells. **(B)** During *Ah* infection, bacteria may escape autophagy and apoptosis of infected cell by affecting Toll like receptor (TLR3) and C-reactive lectin (CLEC4e) mediated signalling. TLR and CLEC4 identify pathogen associated molecular patterns to activate signaling cascades for inflammatory and antimicrobial effects including apoptosis. TLR3 stimulates the TRIF mediated signaling involving TRAF that finally activates MAP kinase (MEK1/2), NF-Κb or Activator protein (AP) pathways. CLEC4e works through ITAM-tyrosine kinase SYK pathway to activate signaling cascade through PLCγ2 for inducing the NFAT/ calcineurin pathway. **(C)** During *Ah* infection, host (immune) cells undergo metabolic reprogramming that may be beneficial for inflammatory host response. The increase in the rate of glycolysis in the host cell degrades glucose into pyruvate through a series of reactions. Expression of glycolysis linked enzymes (Pfk2, Eno1, Ugp2) supports glycolytic flux to form pyruvate. Pyruvate enters into the Citric acid cycle (TCA). Infection mediated metabolic reprogramming leads to functional breaks in the TCA cycle which favour the accumulation of Citrate and Succinate. Downstream metabolite of Citrate (Itaconate) inhibits SDH activity that further increases Succinate. These metabolites increase the mitochondrial reactive oxygen species (mROS) and (Nitric oxide) that have inflammatory effects. Succinate accumulation and redox-environment changes in mitochondria interfere with electron transport chain and stabilise the HIF-1α leading to the expression of MHC Class II and IL-1β. Activated HIF-1α and IL-1β further increase glycolysis and lactate production by increasing Lactate dehydrogenase activity. Fatty acid synthesis increases as a result of Citrate accumulation in cytosol and is required for energy homeostasis and inflammation that may be utilised by the bacteria. In the late infection stage, PPAR signaling gets activated and regulates the ROS mediated apoptosis and anti-inflammatory response. (Single red down arrow-Down regulated protein, Double red down arrow-Protein enriched in controls and not detected/quantified in AH group, Single green up arrow-Upregulated protein, Double green up arrow-Protein enriched in AH group and not detected/quantified in Control group).

Further we propose that *Ah* infection may promote the survival of infected cell by escaping apoptosis or antimicrobial events by interfering with Tlr3 signaling. Immune related protein, Tlr3 was found to be downregulated and another protein involved in the downstream signaling; Dual specificity mitogen-activated kinase kinase 2-like protein (Map2k2b-MEK1/2) was not detected in the infected group (Fig. 5B). TLRs, which recognise pathogen-associated molecular patterns (PAMPs) arising from microorganisms, are one of the most important components of innate or non-specific immunity (20). TLRs can stimulate the TIR-domain-containing adapter-inducing interferon-dependent signaling pathway (TRIF) mediated signaling through Tumor necrosis factor receptor–associated factor (TRAF) that finally activates MAP kinase (MEK1/2), NF-κB or Activator protein (AP) (34). Tlr3 has generally been recognised as endosomal receptor, whereas in *Labeo rohita* and *Cyprinus carpio,* it has been reported as a cell surface receptor capable of TRIF mediated signaling (35). Tlr3, was the first identified antiviral TLR member that has been reported to detect dsRNA of many RNA viruses and activate the TIRF pathway, which produces type I interferon (IFN) and pro-inflammatory cytokines (36). However, Tlr3 has also been observed to respond to bacterial infection in zebrafish, channel catfish, yellow croaker and mice and to respond to Tlr2 microbial ligand peptidoglycan in immature dendritic cells (36). An increased level of Tlr3 at transcriptomic level have been reported in Zebrafish in response to the infection with *Edwardsiella tarda* (37). Our results are similar to those observed in kidney and spleen of blue catfish (38) and kidney of rainbow trout (39) where Tlr3 expression was decreased after infection with gram negative bacteria *E. ictulari* and *Yersinia ruckeri*, respectively. Also, Tlr3 downregulation has been related to immunosuppression in case of severe fever caused by Dabie bandavirus (40). Tlr3 downregulation has been reported as a way to avoid apoptosis during liver cancer (41, 42).

We observed a decreased abundance of Clec4e (Cd207) and Calcineurin in the *Ah* infected group (Fig. 5B). Clec4e protein is a calcium-dependent lectin that acts as a pattern recognition receptor (PRR) of the innate immune system and has been reported to be important for autophagy and antimicrobial responses. Such C-type lectin receptor (CLRs) are known to identify PAMPs and damage associated molecular patterns (DAMPs) and to activate NF-κB signaling for the generation of cytokines and chemokines through the immune-receptor tyrosine-based activation motif (ITAM) and SYK tyrosine kinase pathway (43). Clec4e has been shown to cause signaling through Phospholipase C gamma 2 (PLCγ2) to activate the calcineurin/NFAT pathway (43). Activated Calcineurin stimulates NFAT and NF-κB signaling in T cells, regulates cell growth functions and apoptosis. The NFAT pathway is also involved in the generation of antibodies and the differentiation of B cells (44). Decreased abundance of Clec4e and Calcineurin indicates that *Ah* pathogenesis might involve Clec4e mediated signaling to avoid antimicrobial effects and autophagy of the infected host cell. A combination of Clec4e and Tlr4 agonists has been reported to inhibit the growth of *Mycobacterium tuberculosis* in the lungs of *Mtb-*infected guinea pigs and mice (20). Similar immunotherapeutic approach using Tlr3 can be designed for controlling *Ah* infection.

A few dysregulated proteins belong to Proteosome which is a protease complex involved in hydrolysis of selected proteins in an ATP dependent manner. The proteasome complex has been shown to regulate LPS-induced signal transduction, suggesting that it could be a promising therapeutic target in Gram-negative bacteria. Alpha and beta subunits (Psma1 and Psmb4) of the 20S proteasome complex have been identified as LPS-binding proteins in *Aeromonas salmonicida* infected rainbow trout (45). Further, mapping of upregulated proteins to ribosomal subunits, ribosome biogenesis, amino acyl-tRNA complex and protein processing in ER reveals an increased level of protein synthesis during *Ah* infection. Such biological processes are essential for maintaining homeostasis and promoting host cell immune response against pathogen during infections. Protein synthesis is also necessary for the survival of bacteria inside the host, as reported in case of *Rhodopseudomonas palustris* infection (23).

Further we observed metabolic reprogramming of the host cells as a result of *Ah* infection (Fig. 5C). Proteins related to energy metabolic processes including glycolysis, Kreb’s cycle, and oxidative phosphorylation have showed changes in response to *Ah* infection. Such a reprogramming is more likely an adaptation to fulfil the high energetic and biosynthetic demands in the infected host cell. Among the metabolic enzymes that were altered with *Ah* infection, enzyme 6-Phosphofructo-2-kinase (Pfk2) showed an increased abundance. This enzyme has been reported to play important role in maintaining glycolysis by allosterically regulating the Phosphofructokinase 1 (Pfk1), a rate limiting enzyme of glycolysis (46). Another glycolytic enzyme, Enolase (Eno1) is also upregulated in the AH group. Eno1 converts phosphoglycerate to phosphoenolpyruvate that finally gives pyruvate. Another enzyme involved in carbon metabolism (47); UTP--glucose-1-phosphate uridylyltransferase (Ugp2) was also decreased by 1.7 folds which might be an indication of decreased glycogenesis in support of glycolysis. Ugp2 is also reported to trigger immune responses through P2Y signaling pathway (48). Upregulation of glycolytic enzymes could be a remodelling of the *Ah* infected cell for cell proliferation, similar to what has been shown in cancer cells that rely on high rate of glycolysis (49). Similar findings of glycolytic increase have been reported for mammalian macrophages infected with *Mycobacterium tuberculosis (Mtb)* (50). Increase in the glycolytic enzymes has been observed in kidney derived M1 macrophages of Carp upon activation with bacterial LPS from *Escherichia coli* (51). M1 macrophages undergo metabolic reprogramming from OXPHOS to glycolysis after activation with bacterial LPS alone or in conjunction with IFN-γ (51). In *Mtb* infections, glycolytic reprogramming of M1 macrophages has been associated with two breaks in the tricarboxylic acid cycle (TCA cycle) and suppression of part of the electron transport chain (ETC) in the mitochondria. Breaks in TCA cycle includes the downregulation of two important enzymes i.e., Succinate dehydrogenase (SDH) and isocitrate dehydrogenase (IDH) that favours the build-up of Citrate and Succinate (52, 53).

These reported findings are complementary with our results as we could find higher expression of Acetyl-coenzyme A synthetase (Acss2), Citrate synthase (Cs) and Succinyl ligase (Suclg2) and a downregulation of Isocitrate dehydrogenase (IDH) which has shown enriched expression in the control group. Enzyme Cs catalyses the first reaction of TCA cycle where Oxaloacetate combines with Acetyl-CoA (involving Acss2) to form Citrate (Fig. 5C). The enzyme Succinyl ligase also known as Succinate synthase converts succinyl-CoA to Succinate and free coenzyme A. The downstream metabolite of Citrate (Itaconate) inhibits Succinate dehydrogenase (SDH) activity that again leads to an increase in Succinate (52). Both Citrate and Succinate have been reported to be involved in pro-inflammatory immune functions in macrophages. Citrate can accumulate in the cytosol to play important role in the production of mitochondrial reactive oxygen species (mROS), Nitric oxide (NO) and fatty acid synthesis. mROS can pose antimicrobial effects and activate interleukin 1 β (IL-1β) (54). Intracellular lipid metabolism promotes lipid droplet formation that are stored as energy resource for the cell required for inflammation, cell signaling and homeostasis. They can be utilised by the bacteria as well for promoting infection (55). In the later stages of infection, fatty acid metabolism may stimulate the peroxisome proliferator-activated receptor gamma (PPARγ) signaling which is known for its anti-inflammatory effects through inhibition of proinflammatory cytokines such as IL-1 *β and* TNF-*α* (56). Interestingly, we observed an increased expression of 15-oxoprostaglandin 13-reductase (Ptgr2) during *Ah* infection. Overexpression of Ptgr2 in the cells has been reported to decrease the PPARγ dependent transcription (57). Moreover, Ptgr2 knockdown in LPS stimulated macrophages, resulted in decreased production of pro-inflammatory cytokines (58). Such observations paved a way towards exploring Ptgr2 mediated anti-inflammatory therapy for *Ah* infection. mROS can pose antimicrobial effects and activate interleukin 1 β (IL-1β) (54).

Succinate accumulation also leads to the generation mROS and NO and can stabilise hypoxia-inducible factor 1-alpha (HIF-1α). Activated HIF-1α further promotes the glycolytic pathway and lactate production and causes inflammation by positively regulating the glycolytic genes and IL-1β expression (52, 59). In our study, Lactate dehydrogenase enzyme (Ldha) has been found to show enriched expression in the *Ah* infected group (AH group). During this reprogramming, mitochondrial electron transport chain (ETC) is highly disturbed (52). In our study, we observed dysregulation of many proteins involved in ETC and OXPHOS. Dysregulated proteins include NADH dehydrogenase (Ndufa, Ndufb, Ndufv2, Ndufs8), ATP synthase-coupling factor 6 (ATP5J), ATP synthase protein 8 (ATP8), Mt-co1(Cytochrome c oxidase subunit 1), and Inorganic diphosphatase (Ppa2) (Fig. 5C). Such a reprogramming has been profoundly observed in LPS activated dendritic cells and macrophages, activated effector T cells, activated natural killer cells and activated B cells (60). Remodelling of these metabolic processes enables the cells to produce sufficient ATP for performing cellular functions which may be inflammatory cytokine production, phagocytosis or antigen presentation (60). Based on these findings and our observations, we hypothesise that *Ah* pathogenesis in *Labeo rohita* involves metabolic reprogramming that supports several inflammatory immune functions which is a hallmark for most immune cells.

## CONCLUSIONS

We analysed the proteome level dynamics in liver tissue of *L. rohita* as a result of *Ah* infection. Our proteomics analysis provided new insights into the proteins involved and processes underlying *Ah* pathogenesis in *L. rohita.* We observed that during *Ah* infection, several host proteins and metabolic pathways were altered affecting important biological processes like antioxidative potential, lysosomal killing and apoptosis. We also observed remodelling of important energy related processes such as glycolysis and Krebs cycle which might be useful in modulating inflammatory response during *Ah* infection in *L. rohita*. Our findings have cleared the way for more research into the involvement of Toll-like receptors (Tlr3), C-type lectins (Clec4e), and metabolic enzymes in *Ah* pathogenesis. Collectively, our data provided new mechanistic insights into *Ah* infection that can help in characterisation of host-pathogen interactions and aid in the selection and prioritisation of proteins to be used in host directed immunotherapy against *Ah* infection.

## MATERIALS AND METHODS

### Overall experimental design

This study aimed at proteomic profiling of liver tissue of *Ah* infected *Labeo rohita*. Fishes were challenged with *Ah* and sampled such that there was a total of 12 samples including six each for Control and AH group (*Ah* infected). Following sample collection, discovery based proteomic analysis was performed by taking four samples for each group (four each of Control and AH group). The data was analyzed using MaxQuant software followed by statistical analysis in Metaboanalyst tool to identify the differentially expressed proteins (DEPs) in liver tissue. Gene ontology (GO) analysis was performed to obtain an overview of functional annotation of significant proteins and dysregulated metabolic pathways. The protein abundance changes for a panel of differentially expressed proteins was validated using SRM approach where eleven samples (6 Control and 5 AH) were analyzed. Further, detailed analysis was done to understand the molecular mechanism of *Ah* infection.

### Bacterial collection and identification

In the study, *Aeromonas hydrophila* strain was isolated from the kidney tissue of naturally co-infected *Labeo rohita* (NCBI Accession no. MT374248). Briefly, kidney tissue was streaked onto tryptic soy agar (TSA, Himedia) and incubated at 28 °C for 24 hours. Representative colonies were isolated and re-streaked on fresh TSA medium until purity was attained. Pure cultures of the isolated bacteria were subjected to morphological analysis, and the taxonomy of the isolates was determined following the 16S rRNA gene using the gene sequence universal primers; forward primer 27F 5’-AGAGTTTGATCCTGGCTCAG-3’ and reverse primer 1492R 5’-GGTTACCTTGTTACGACTT-3’ (Chromous Biotech Ltd., Bengaluru, India) (61) (Fig. 1A). The strain was stored at −70 °C in 30% glycerol stock until use for the challenge study.

To detect the virulence of strain, *Ah* (MT374248) was cultured in Brain Heart Infusion broth (BHI) (Himedia) at 28 °C and kept in incubator at 150 rpm for 18 hours. For confirmation of *Ah*, a loop of bacteria was streaked onto Aeromonas Isolation Medium HiVegTM Base (Himedia) and polymerase chain reaction (PCR) was conducted using target segments of primers; forward: 16S rDNA1 451–473 5′-GAAAGGTTGATGCCTAATACGTA-3′ and reverse:16S DNA2 1115–1135 5′-CGTGCTGGCAACAAAGGACAG-3′ with an expected product length of 685 bp (62). The amplifications were performed in thermal cycler (Bio-Rad, USA) in which 30 PCR cycles were run under the following conditions; denaturation at 94 °C for 2 min, primer annealing at 56 °C for 2 min, and DNA extension at 72 °C for 2 min in each cycle. A negative control with all the reaction components except template DNA and a positive control (extracted genomic DNA, from *Ah* MTCC culture, 1739T-Chandigarh, India) were considered. Ten microliters of PCR products were electrophoresed in a 1% agarose gel containing ethidium bromide at 100 V for 1 hour, visualized on an ultraviolet (UV) transilluminator (Bio-Rad) (Fig. S5).

### Collection of fish and maintenance for the experiment

For expression study, six-month-old fish (N=150, average weight 70±10 g) were collected from a local fish farm (Pen Raigad District, Maharashtra) and brought to wet lab facility (ICAR-CIFE, Mumbai). Fishes were equally distributed and acclimated in three circular fibre tanks and fed 2 % body weight. Fishes were maintained at a temperature of 26-28 °C with proper aeration, and removal of faeces on daily basis. All the fishes were observed for external clinical signs and randomly few fishes from three tanks sacrificed to observe pathogen in the fish using PCR.

### *Aeromonas hydrophila* challenge and sampling

For challenge test, LD50 dose was determined as per the protocol described by Siriyappagouder et al. (63) and found to be 1.5 ×10^8^ CFU for the isolated strain. The bacteria were cultured till 18 hours, and LD50 dose of 1.5 ×10^8^ CFU suspension was prepared after washing with PBS. The fish were pooled and acclimatized, starved for 2 days and then approximately 1.5×10^8^ bacterial cells in PBS or the same volume of PBS solution (Control) were intraperitoneally inoculated into 36 fish (18 each for control and challenged). The challenged and control groups were separately maintained in six crates (6 fish each) of a 100-liter capacity high density polyethylene plastic crate at 26-28 °C. The presence of external signs of hemorrhage were observed in the challenged fish groups post-infection. Expectedly, these signs were not detected in the control group (Fig. S6). After 48 hours, fish were euthanized and liver samples were collected. Collected samples were stored at −80 C till further use. For proteomic analysis, tissue from three fishes were pooled into one resulting in a total of six samples each for control and challenged group labelled as Liv-C1 to C6 and Liv-AH1 to AH6, respectively.

### Tissue lysis and protein extraction

Tissue lysates were prepared using SDS containing lysis buffer (5% SDS, 100mM Tris/HCl pH 8.5 (adjusted with phosphoric acid). The tissue was weighed (40-50 mg) and rinsed in a 1X phosphate buffered saline (PBS) solution (2-3 times to remove any blood). After washing the tissue, 250 µl of lysis buffer was added to the tissue along with 5 µl of Protease inhibitor cocktail (50X stock, Sigma-Catalogue no. 11873580001) and incubated on ice for 30 min. Sonication was performed for 2 min with an amplitude of 40% with 5 sec pulse on and 5 sec off. Centrifugation was done to remove the debris and clear supernatant was collected.

### Protein quantification and digestion

Tissue lysates were processed for quantification of proteins using the BCA assay (Thermo, Ref. 23227) with Bovine serum albumin as the standard protein. After quantification, 30 µg protein was taken for digestion using filter assisted sample preparation (FASP) based digestion method. In brief, the protein was first reduced using TCEP solution with a final concentration of 20 mM. Initial volume at this step was kept as 30 µl (volume made up using 1X lysis buffer). Reduced sample was loaded onto a 30 KDa filter (Catalogue no. MRCF0R030-Merck Millipore) to proceed for FASP based digestion. Following the required steps of alkylation and washing, trypsin mixed in digestion buffer (50 mM Ammonium Bicarbonate) was added in 1:30 ratio for enzyme to protein. Samples were incubated in a wet chamber at 37°C for 16 hours. Digested peptides were eluted in a fresh collection tube, dried and stored until further processing. Before mass spectrometry, samples were cleaned using C18 stage tips (Empore™ SPE Disks matrix active group C18, diam. 47 mm, catalogue no. 66883-U-Merck).

### Liquid chromatography tandem mass spectrometry in data dependent acquisition mode

For all the samples, peptides were quantified using Scopes method (64). After quantification, one µg of peptide sample was loaded on the column and LC-MS/MS was performed. All samples (4 for each Control and AH group) were run with an LC gradient of 120 min. Data was acquired using an Orbitrap-Fusion Tribrid mass-spectrometer connected to an Easy-nLC nano-flow liquid chromatography 1200 system. Peptide sample was loaded onto the pre-analytical column (100 µm x 2 cm, nanoViper C18, 5 µm, 100A; Thermo Fisher Scientific) at a flow rate of 5 µl/min. Peptides were resolved on analytical column (75 μm × 50 cm, 3 μm particle, and 100 Å pore size; Thermo Fisher Scientific) at a flow rate of 300 nl/min over 120 min gradient in solvent B (80% Acetonitrile with 0.1% Formic acid (FA). The Orbitrap mass analyzer was used to perform mass spectrometric acquisition in data dependent acquisition (DDA) mode in the full scan range of 375-1700 m/z with a mass resolution of 60,000. With a dynamic exclusion time of 40 seconds, the mass window was set at 10 ppm. All MS/MS spectra were obtained using the HCD method (High Energy Collision Dissociation) for fragmentation at MS1 and MS2 level, the AGC target was set at 400000 and 10000, respectively. A lock mass of 445.12003 m/z was used for positive internal calibration.

### Protein identification and quantification using label-free quantification approach

The raw mass spectrometry data was analysed using MaxQuant (v1.6.6.0) software against UniProt protein database for *Labeo rohita* (ProteomeID-UP000290572, Taxonomy ID-84645, downloaded on 18.06.2021) using the in-built search engine, Andromeda. All the raw files were analyzed together using Label-Free-Quantification (LFQ) parameters. Label type was set to standard with a multiplicity of 1, and the match between run option was checked. Orbitrap fusion mode was selected as instrument, and trypsin as a protease was chosen. A total of two missed cleavages were permitted. Carbamidomethylation at Cysteine (+57.021464 Da) was chosen for fixed modification, and oxidation at Methionine (+15.994915 Da) was chosen for variable modification. A false discovery rate of 1% was specified for both proteins and peptides. Reverse was chosen as the decoy mode option, and proteins were recognised solely by their unique peptide. The LFQ intensities obtained for each sample were considered for quantitative analysis (Table S1).

### Statistical analysis using MetaboAnalyst

Statistical analysis was performed taking MaxQuant analyzed files in Metaboanalyst software (65). One of the samples (Liv-AH1) was discarded due to poor mass spectrometry results (Fig. S1A) The proteomic data of the remaining seven samples was considered for further analysis including four Control samples; Liv-C1, 2, 3, 4 and three *Ah* infected samples (AH group) labelled as Liv-AH2, 3, 4. The K-Nearest Neighbor (KNN) algorithm was used to impute the missing values of proteins with abundance values in more than 75% of each group, which were then used in differential protein expression analysis. The data was log_10_ transformed before further statistical analysis. The significant differentially expressed proteins were identified using a two-sample t-test (Welch t-test) with a p-value threshold of 0.05. Proteins with minimum fold change value of 1.5 were regarded as significant differentially expressed proteins (DEPs) among all t-test passed proteins. Variable importance in projection (VIP) score was obtained through Partial Least Squares Discriminant Analysis (PLS-DA) which is a supervised method. As a weighted sum of squares of the PLS weight, VIP indicates the importance of the variable to the whole model. Heatmap representing the expression of top DEPs was also obtained from Metaboanalyst analysis. Volcano plots were plotted using online tool VolcanoseR (66).

### Gene ontology, Pathway and protein-protein interaction enrichment analysis

Selected dysregulated proteins obtained from LFQ data were taken forward for functional annotation and biological pathway analysis. Gene names of dysregulated proteins were retrieved from EggNOG resource (67) on the basis of ortholog annotation and from literature (as the gene names for *L. rohita* are not updated yet in the available databases). The protein-protein interaction (PPI) enrichment analysis and visualization of their involvement in respective biological pathways (KEGG and Reactome) was performed using STRING tool version 11.5 (68) taking genes of significantly upregulated and downregulated proteins as input (Table S2). *Danio rerio* was selected as reference organism as these databases are not yet updated for *Labeo rohita*. Furthermore, biological functional pathway enrichment of different Gene Ontologies (GO), KEGG, Reactome and Panther Pathway and PPI were done using Metascape tool (69). In Metascape, custom analysis of DEPs, control and disease enriched proteins were performed selecting *Danio rerio* as reference organism, considering all of the terms and categories for annotation and membership (Table S2). Minimum overlap of 3, p-value cut off of 0.05, minimum enrichment 1.5 were taken as parameters for pathway and process enrichment and physical core database was selected for protein-protein interaction enrichment analysis.

### Targeted proteomic validation of dysregulated proteins using SRM approach

For acquiring the targeted proteomic data using selected/ multiple reaction monitoring (SRM/MRM) approach, TSQ Altis mass spectrometer (ThermoFisher Scientific, USA) coupled to an HPLC-Dionex Ultimate 3000 system (ThermoFisher Scientific, USA) was used.

In order to separate the peptides, a Hypersil Gold C18 column (1.9 μm, 100 x 2.1 mm, ThermoFisher Scientific, USA) was used. The flow rate was maintained as 0.45 ml/min for 10 min. Solutions of 0.1% FA and 80% ACN in 0.1% FA were taken as buffer A and B, respectively in the binary buffer system. The gradient used for chromatographic separation of the peptides was as follows; 2-45% buffer B for first 6 min, 45-95% buffer B for 0.5 min, 95% buffer B for 0.5 min, 95%-2% buffer B for 0.5 min, and 2% buffer B for 2.5 min.

We started with a list of 30 proteins (16 upregulated and 17 downregulated) corresponding to 244 peptides and ~4000 transitions. The Skyline software, version 20.1.1.196 (70) was used to prepare the transition lists to be fed into the system. The criteria included for miss cleavage was 0, for precursor charges +2, +3, and product charge was +1 with ‘y’ ion transitions (from ion 2 to last ion −1). A background proteome consisting of UniProt protein database for *Labeo rohita* (ProteomeID-UP000290572, Taxanomy ID-84645, downloaded on 18.06.2021) was used. Unique peptides previously identified in the discovery data were only selected for the targeted experiment. The collision energy values used in the experiment were as determined by Skyline software. One μg of peptides from each sample were injected and run against the target list. Initial optimization was done using pooled peptide samples which were run against nine transition lists each with 400-450 transitions per method. Consequently, the list was refined based on consistency of the spectral data. Before injecting to mass spectrometer, all the samples were spiked in with equal amount or heavy labelled synthetic peptide DIFTGLIGPMK (C-terminus lysine labelled). Synthetic peptide was added to monitor the consistency of the mass spectrometric run, for which 18 transitions were added to the final transition list. For the final SRM run, the data was acquired for all the samples (five AH and six Control) using two transition lists (two SRM methods) consisting of 617 transitions and 91 peptides corresponding to 27 proteins resulting in 22 raw files (Table S3).

### Targeted proteomics data analysis

After data acquisition, all the downstream data analysis was performed in Skyline software (70). MSMS (.msms) file obtained from the Maxquant analysis was utilised for preparing the DDA spectral library for analysing S/MRM data. The result files (.raw) were imported to the Skyline software and assigned to respective conditions as Control and AH. Each peptide was manually annotated based on peak shape and retention time alignment with other replicates of the same peptide and library match (dot product measurement). The peptides that didn’t adhere well to the retention time alignment were deleted after manual annotation to refine the data. Statistical analysis was performed using MSstats external tool inbuilt in Skyline in which certain peptides were found after the fold change analysis (cut off 1.5) with significant p-value (0.05). Result reports for all the peptides containing their peak area values were exported in .csv format to carry out the further analysis (Table S3). Further data analysis was done for proteins with two or more significant peptides and violin plots for peptide wise intensities (Table S3) were created using an online tool BoxPlotR (71).

## DATA AVAILABILITY

The protein database (.FASTA) and raw mass spectrometry data (.raw) have been deposited to the ProteomeXchange Consortium via the PRIDE partner repository. All result output files for protein identification are also submitted in text (.text) format along with the parameter file. Also, the spectral library (.blib) generated using the discovery data for analysing the targeted data is uploaded. The identifier PXD029421 can be used to retrieve all of the data. (Reviewer account details: Username, Password: lI191yU0).

The transition lists, skyline documents, and all SRM raw (.raw) data for the Selected reaction monitoring (SRM) experiment have been submitted to Panorama public that can be accessed through the given link https://panoramaweb.org/rohuliverproteomicsah.url (Reviewer account details: Password: OKRYbqkf).

## AUTHOR’S CONTRIBUTIONS

Concept and design: M.N., S.S. and M.G.

Maintenance and sampling: N.P., M.N. and M.G.

Method development and Data acquisition: M.N. and S.S

Data analysis and Interpretation: M.N., N.P., B.G. and U.S.

Writing and review: M.N., N.P., B.G., U.S., M.G. and S.S

## Abbreviations

Aars: Alanine--tRNA ligase OS
Atp2b1: Calcium-transporting ATPase
Atp6v1a: H(+)-transporting two-sector ATPase
Atp6voc: V-type proton ATPase proteolipid subunit
Atp8: ATP synthase protein 8 OS
Cpt1: Carnitine O-palmitoyltransferase OS
Cyp1a: Unspecific monooxygenase OS
Cyp2f2: Cytochrome P450 2F2-like protein OS
Cyp2g1: Cytochrome P450 2G1-like protein
Cyp3a: Cytochrome P450 3A30-like protein OS
Eprs: Glutamyl-tRNA synthetase OS
Fabp: Fatty acid-binding brain-like protein OS
Hsp90aa1: Heat shock HSP 90-alpha OS
Hspa8: Heat shock cognate 71 kDa OS
Mogs: Mannosyl-oligosaccharide glucosidase-like protein OS
Ndufa12: NADH dehydrogenase [ubiquinone] 1 alpha subcomplex subunit 12
Ndufa2: Complex I-B8 OS
Ndufa6: Complex I-B14
Ndufb10: Complex I-PDSW OS
Ndufb6: Complex I-B17
Ndufs8: Complex I-23kD
Ndufv2: NADH dehydrogenase [ubiquinone] flavoprotein 2, mitochondrial
Psma1: Proteasome subunit alpha type-1
Psma5: Proteasome subunit alpha type OS
Psma8: Proteasome subunit alpha type OS
Psmb4: Proteasome subunit beta
Psmd3: 26S proteasome non-ATPase regulatory subunit 3 OS
Psmd6: 26S proteasome non-ATPase regulatory subunit 6 OS
Qars: Glutamine--tRNA ligase OS
Rpl13: 60S ribosomal protein L13 OS
Rpl7a: 60S ribosomal protein L7a OS
Rplp2: 60S acidic ribosomal protein P2 OS
Rps6: 40S ribosomal protein S6 OS
Rrbp1: Ribosome-binding 1-like isoform X1 OS
Scp2: Acetyl-CoA C-myristoyltransferase OS
Sec31a: Transport Sec31A-like isoform X5 OS
TCEP: Tris(2-carboxyethyl)phosphine
Tlr4: Toll-like receptor 4

## ACKNOWLEDGEMENTS

This work was supported by Department of Biotechnology (BT/PR15285/AAQ/3/753/2015) Govt. of India to S.S and M.G. M.N was supported by University Grants Commission (UGC). We acknowledge ICAR-Central Institute of Fisheries Education, Mumbai for supporting this work. We acknowledge MASS-FIITB at IIT Bombay supported by the Department of Biotechnology (BT/PR13114/INF/22/206/2015) for Mass-spectrometric data acquisition.

## SUPPLEMENTAL INFORMATION

This article contains six supplementary figures (Figure S1 to S6) and three supplementary Tables (Table S1 to S3) (.xls).

